# High pathogenicity avian influenza (H5N1) in Northern Gannets: Global spread, clinical signs, and demographic consequences

**DOI:** 10.1101/2023.05.01.538918

**Authors:** Jude V Lane, Jana WE Jeglinski, Stephanie Avery-Gomm, Elmar Ballstaedt, Ashley C Banyard, Tatsiana Barychka, Ian H Brown, Brigitte Brugger, Tori V Burt, Noah Careen, Johan HF Castenschiold, Signe Christensen-Dalsgaard, Shannon Clifford, Sydney M Collins, Emma Cunningham, Jóhannis Danielsen, Francis Daunt, Kyle JN d’Entremont, Parker Doiron, Steven Duffy, Matthew D English, Marco Falchieri, Jolene Giacinti, Britt Gjerset, Silje Granstad, David Grémillet, Magella Guillemette, Gunnar T Hallgrímsson, Keith C Hamer, Sjúrður Hammer, Katherine Harrison, Justin D Hart, Ciaran Hatsell, Richard Humpidge, Joe James, Audrey Jenkinson, Mark Jessopp, Megan EB Jones, Stéphane Lair, Thomas Lewis, Alexandra A Malinowska, Aly McCluskie, Gretchen McPhail, Børge Moe, William A Montevecchi, Greg Morgan, Caroline Nichol, Craig Nisbet, Bergur Olsen, Jennifer Provencher, Pascal Provost, Alex Purdie, Jean-François Rail, Greg Robertson, Yannick Seyer, Maggie Sheddan, Catherine Soos, Nia Stephens, Hallvard Strøm, Vilhjálmur Svansson, T David Tierney, Glen Tyler, Tom Wade, Sarah Wanless, Christopher RE Ward, Sabina Wilhelm, Saskia Wischnewski, Lucy J Wright, Bernie Zonfrillo, Jason Matthiopoulos, Stephen C Votier

**Affiliations:** RSPB Centre for Conservation Science, Sandy, UK; School of Biodiversity, One Health and Veterinary Medicine, University of Glasgow, Scotland, UK; Wildlife and Landscape Science Directorate, Science & Technology Branch, Environment and Climate Change Canada; Verein Jordsand zum Schutz der Seevögel und der Natur e. V.; International Reference Laboratory for Avian Influenza-Weybridge, Animal and Plant Health Agency, Addlestone, Surrey UK; Icelandic Food and Veterinary Authority, Iceland; Psychology Department, Memorial University of Newfoundland and Labrador, Canada; Aarhus University, Dept. of Ecoscience, Frederiksborgvej 399, 4000 Roskilde, Denmark; Norwegian Institute for Nature Research (NINA), PO Box 5685 Torgard, 7485 Trondheim, Norway; Centre for Immunity, Infection and Evolution, Institute of Evolutionary Biology, School of Biology, University of Edinburgh, Scotland, UK; Seabird Ecology Department, Faroe Marine Research Institute, Nóatún 1, FO-100 Tórshavn, Faroe Islands; UK Centre for Ecology & Hydrology, Bush Estate, Penicuik, Scotland, UK; Canadian Wildlife Service, Environment and Climate Change Canada; Norwegian Veterinary Institute, PO Box 64, N-1431 Ås, Norway; CEFE, Univ Montpellier, CNRS, EPHE, IRD, Montpellier, France; FitzPatrick Institute, DST/NRF Excellence Centre at the University of Cape Town, Rondebosch 7701, South Africa; Department of Biology, Chemistry and Geography, Université du Québec à Rimouski, Rimouski, Québec, Canada; Department of Life and Environmental Sciences, University of Iceland, Reykjavik, Iceland; School of Biology, University of Leeds, UK; Faroese Environment Agency, Traðargøta 38, FO-165 Argir, Faroe Islands; University of the Faroe Islands, J. C. Svabos gøta 14, FO-100 Tórshavn, Faroe Islands; Department of Agriculture, Food and the Marine (DAFM), Agriculture House, Kildare Street, Dublin, Ireland; Alderney Wildlife Trust, Channel Islands; National Trust for Scotland, Hermiston Quay, Edinburgh, Scotland, UK; RSPB Scotland, UK; School of Biological, Earth & Environmental Sciences, University College Cork, Ireland; University of Prince Edward Island, Canada; Centre québécois sur la santé des animaux sauvages, Canadian Wildlife Health Cooperative, Faculté de médecine vétérinaire, Université de Montréal, St. Hyacinthe, Québec, Canada; RSPB Hoy, Stromness, UK; RSPB Ramsey Island, St Davids, Pembrokeshire UK; School of Geosciences, University of Edinburgh, UK; Ligue pour la Protection des Oiseaux, Réserve Naturelle Nationale des Sept-Iles, Pleumeur Bodou, France; Scottish Seabird Centre, North Berwick, UK; Norwegian Polar Institute, Fram Centre, 9296 Tromsø, Norway; Institute for Experimental Pathology, Biomedical Center, University of Iceland, Keldur, Iceland; Science and Research Directorate, National Parks and Wildlife Service, 90 King Street North, Dublin 7, D07 N7CV, Ireland; NatureScot, Great Glen House, Leachkin Road, Inverness IV3 8NW, UK; Lyell Centre, Institute for Life and Earth Sciences, Heriot-Watt University, Edinburgh, UK

**Keywords:** HPAIV, avian flu, virus outbreak, seabirds, immunity

## Abstract

During 2021-22 High Pathogenicity Avian Influenza (HPAI) killed thousands of wild birds across Europe and North America, suggesting a change in infection dynamics and a shift to new hosts, including seabirds. Northern Gannets (*Morus bassanus*) appeared especially severely impacted, but limited understanding of how the virus spread across the metapopulation, or the demographic consequences of mass mortality limit our understanding of its severity. Accordingly, we collate information on HPAIV outbreaks across most North Atlantic gannet colonies and for the largest colony (Bass Rock, UK), provide impacts on population size, breeding success, adult survival, and preliminary results on serology. Unusually high numbers of dead gannets were first noted in Iceland during April 2022. Outbreaks in May occurred in many Scottish colonies, followed by colonies in Canada, Germany and Norway. By the end of June, outbreaks had occurred in five Canadian colonies and in the Channel Islands. Outbreaks in 12 UK and Ireland colonies appeared to follow a clockwise pattern with the last infected colonies recorded in late August/September. Unusually high mortality was recorded at 40 colonies (75% of global total colonies). Dead birds testing positive for HPAIV H5N1 were associated with 58% of these colonies. At Bass Rock, the number of occupied sites decreased by at least 71%, breeding success declined by ∼66% compared to the long-term UK mean and adult survival between 2021 and 2022 was 42% lower than the preceding 10-year average. Serological investigation detected antibodies specific to H5 in apparently healthy birds indicating that some gannets recover from HPAIV infection. Further, most of these recovered birds had black irises, suggestive of a phenotypic indicator of previous infection. Untangling the impacts of HPAIV infection from other key pressures faced by seabirds is key to establishing effective conservation strategies for threatened seabird populations, HPAIV being a novel and pandemic threat.

## 1. Background

High Pathogenicity Avian Influenza Virus (HPAIV) H5Nx has negatively impacted wild and domestic bird populations globally for decades (Nuñez and Ross. 2019). However, the current global panzootic of H5Nx has seen shifts in both the seasonality of outbreaks and the species affected (EFSA, 2023). H5Nx (A/goose/Guangdong/1/1996 (Gs/GD) H5N1) was first detected in 1996 on a domestic goose farm in Guangdong Province, China (Xu et al. 1999). This goose Guangdong lineage (Gs/Gd) has since caused significant outbreaks in a variety of bird populations and has also raised concerns about the potential zoonotic consequences for humans (Wan 2012, EFSA 2023). Genetic reassortment has led to the emergence and evolution of multiple subtypes and genotypes of this group of high pathogenicity viruses on a global scale, potentially with different epidemiological properties, especially with respect to host range in wild birds (Monne et al. 2014, Falchieri et al. 2022). The mechanism of viral transmission is likely a combination of infected wild bird migration and the global domestic poultry trade or their products (Blagodatski et al. 2021, Ramey et al. 2022).

Low Pathogenicity Avian Influenza Virus (LPAIV) is widely circulating in wild aquatic birds; *Anseriformes* (waterfowl) and *Charadriiformes* (shorebirds) are known to act as reservoirs (Venkatesh et al. 2018) however, we know little about the recent emergence, spread and impact of HPAIVs in aquatic birds, including seabirds (Burggraff et al. 2014, Falchieri et al. 2022, Boulinier. 2023, Roberts et al. 2023). HPAIVs do not evolve within wild bird populations but once they have spilled into wild populations, are transmitted via infected saliva, nasal secretion and faeces but shedding methods differ between species and are not well understood (Arnal et al. 2014, Caliendo et al. 2020).

The winter of 2021/2022 saw a record number of confirmed cases of HPAIV H5N1 in poultry, captive and wild birds across Europe (EFSA 2023). HPAIV H5N1 was first detected in UK breeding seabirds in July 2021 when Great Skuas (*Stercorarius skua*) on Fair Isle, Scotland tested positive (Banyard et al. 2022). The first case of H5N1 detected in North American seabirds was a Great Black-backed Gull (*Larus marinus*) in Newfoundland and Labrador, Canada in November 2021, with phylogenetic analyses revealing the virus was of the European H5N1 lineage (Caliendo et al. 2022). In early April 2022, Common Eider (*Somateria mollissima*) was the first seabird species to test positive for HPAIV in the UK that year, followed in late April by Great Skua (Falchieri et al. 2022). Then followed an unprecedented epidemic in seabirds across the North Atlantic, with Northern Gannets (*Morus bassanus;* hereafter gannet), previously unknown to have been impacted by H5Nx, being severely impacted (Cunningham et al. 2022).

Gannets breed in 53 colonies of various sizes (<10 to >60,000 breeding pairs and non-breeding immatures) on sea cliffs, stacks, and islands across both sides of the North Atlantic from Russia to north-eastern North America (d’Entremont et al. 2022a, Jeglinski et al. 2023). During the breeding season, gannets are medium-range foragers capable of travelling more than 1000 km to find food (Hamer et al. 2007). Moreover, immature gannets travel greater distances than adults and prospect other colonies (Votier et al. 2011, Votier et al. 2017, Grecian et al. 2018). During the non-breeding period, gannets are migratory with birds from Iceland and the eastern Atlantic occupying marine wintering grounds in UK waters, Iberia, with the majority wintering off the coast of West Africa (Veron and Lawlor 2009, Fort et al. 2012, Furness et al. 2018, Deakin et al. 2019). Birds from the western Atlantic primarily winter along the coasts of the eastern United States of America south to the Gulf of Mexico, although some also winter off the coast of West Africa (Fifield et al. 2014). Considering the oral-faecal spread of avian influenza viruses, opportunities for spread between gannets are most likely at the colony but may also occur at foraging grounds and wintering areas by birds in the early stages of infection (Weber and Stilianakis 2007).

Globally, gannets are classified as Least Concern by the International Union for Conservation of Nature (IUCN) due to their wide distribution and growing populations in Europe and North America (IUCN 2023). The European population comprises 75-94% of the global population with 55.6% breeding in the UK (IUCN, 2023). The Bass Rock, Scotland (56° 6′ N, 2° 36′ W) is the world’s largest gannet colony with an estimated 75,259 apparently occupied sites (AOSs) in 2014 (Murray et al. 2015).

Understanding virus spread and infection outcome is essential to fully understand how the HPAIV outbreak impacted gannets and other seabirds. Here, we provide the first comprehensive assessment of the spatio-temporal occurrence of HPAIV outbreaks at most gannet colonies across their North Atlantic breeding range. Moreover, to better understand HPAIV transmission, immunity, and the potential for population recovery, we present detailed results from the largest gannet colony at Bass Rock, Scotland. We quantify how the 2022 HPAIV outbreak influenced adult survival and breeding success and present a non-invasive method that has the potential to determine exposure status based on iris colour.

## 2. Methods

### a. Global context: HPAIV spread across the North Atlantic gannet metapopulation

We collated and mapped data on the timing of unusually high gannet mortalities (the earliest observations notable to fieldworkers familiar with their sites) at existing colonies across the global metapopulation as defined by Jeglinski et al. 2023. Colonies in Norway, Iceland and some Irish colonies were not monitored directly, but instead we gathered information on dead gannet sightings reported to the Norwegian Species Observation System (www.artsobservasjoner.no), the Icelandic Food and Veterinary Authority and to the Department of Agriculture, Food and the Marine’s (DAFM) Avian Check App (https://aviancheck.apps.services.agriculture.gov.ie/), and we associated these observations with the nearest gannet breeding colony. We also collated information on positive HPAIV tests associated with gannet colonies, based on data from the national testing laboratories for the relevant countries.

### b. Case Study: Bass Rock

#### I. Health and safety and biosecurity

Strict biosecurity and health and safety measures were followed to ensure the safety of birds and field workers. During handling our Personal Protection Equipment (PPE) comprised coveralls, face masks, goggles, disposable aprons, and gloves. Safe4 disinfectant was used for disinfecting equipment and footwear (see Supplementary Online Material S1).

#### II. Impact of HPAIV on apparently occupied sites, breeding success and adult survival

##### Apparently occupied sites

A DJI Matrice 300 RTK unmanned aircraft system fitted with a DJI-Zenmuse L1 LiDAR and photogrammetry sensor was flown over the Bass Rock between 15:07-15:19 on 30^th^ June to count live and dead birds. All flights were conducted from the southern tip of the island with a Real-Time-Kinematic (RTK) base station, in good light with light winds (<5ms^-1^) enabling a flight speed of 4 ms^-1^, with image sidelap of 70% and endlap of 80%. The resulting 102 images (captured with 0.001 of a second shutter speed and auto ISO) collected at an altitude of 100 m above ground level, were processed through Agisoft Metashape (Agisoft LLC, St Petersburg, Russia) to produce an orthomosaic of the Bass Rock with a ground sampling distance of approximately 3 cm (see Supplementary Online Material S2). The composite image was loaded into DotDotGoose version 1.5.3 (<u>DotDotGoose</u> (amnh.org)) to allow manual counting of birds on the colony. White birds were presumed to be adults but could not be distinguished from 4-5-year-old immatures. Birds were considered dead based on spread wings or contorted body shape, or alive if their posture was apparently natural or too indistinct to see.

##### Breeding success

We monitored 93 active nests in two study sites, during 14 visits between 15^th^ June and 14^th^ August 2022. All nests had an egg on the first visit, those with a chick on 14^th^ August were considered successful.

##### Adult survival

Visual searches for 370 colour-ringed adults (marked during 2010-2021) took place weekly (total of 12 days) from 15^th^ June until 30^th^ July 2022. Nest sites of colour-ringed birds were repeatedly scanned from a distance of between ∼5-30m and the ring sequence of each bird recorded during a total of ∼11 person-observation hours each day.

We constructed annual encounter histories for each marked bird using resighting data from July 2022 and from visits made in July 2011-2021. A goodness-of-fit test (GOF) showed that a fully time-dependent (both survival (ϕ) and resighting (*p*) probabilities vary with time) Cormack-Jolly-Seber (CJS) model did not fit the data well (GOF: χ^2^ _34_ = 73.33, P = <0.01) with evidence of trap dependence (TEST2.CT; z = -6.1484, two-sided test, p <0.01) but no evidence for transience (TEST3.SR; z = -1.9044, two-sided test, p = 0.056). After accounting for trap-dependence a variance inflation factor (c) of 1.212 was estimated by U-CARE (Choquet et al. 2009). Therefore we set c = 1.212 to account for the over-dispersion in the data and a two-stage TSM structure was applied to model re-sightings.

Models were specified in MARK (Version 9.0, White and Burnham 1999) with the candidate model set (n=4) built so that the survival and resighting probability parameters could vary with year (*t*) or remain constant over time (*c*).

#### III. Serology and iris colour

During September 2022 we caught 19 apparently healthy chick-rearing adults and took ∼1 ml of blood from the tarsal vein (under licence from the UK Home Office; Project licence number PEAE7342F). Sampling effort focused on catching equal numbers of birds with healthy and abnormally black irises, seen for the first time during the outbreak. Birds were caught from seven distinct locations to minimise potential bias in virus exposure between clusters of nests. Where possible, birds with chicks were caught preferentially to guarantee that they had been present throughout the HPAIV outbreak. Birds without chicks were caught if they appeared to be holding a territory.

We took external cloacal swabs from 18 of the 19 birds to test for any possible asymptomatic HPAI infection. Blood and cloacal swabs were stored in a cool bag with ice blocks in the field, then stored at ∼4°C before being transported directly to the UK reference laboratory for avian influenza at the Animal and Plant Health Agency (APHA). Blood samples were tested for an indication of previous infection using an hemagglutination inhibition assay to detect antibodies to H5 avian influenza virus (clade 2.3.4.4b) using a viral antigen homologous to the outbreak virus. Swabs were first tested for influenza A virus nucleic acid following RNA extraction using a matrix (M) gene-specific real-time reverse-transcriptase polymerase chain reaction (rRT-PCR) assay (Nagy et al. 2021) and a HPAIV specific H5 PCR assay (James et al. 2022). Unless already ringed, birds were fitted with a metal British Trust for Ornithology (BTO) ring and a blue plastic darvic ring engraved with a unique alphanumeric code to allow future identification.

A Fisher’s exact test was used to determine the associations between iris colour and exposure status. Statistics were performed using R 4.1.1 (R Development Core Team 2007).

## 3. Results

### a. Global Context: HPAIV spread across the North Atlantic gannet metapopulation

We gathered evidence of HPAI occurrence for 41 colonies. Unusually large numbers of dead gannets were detected at 40 of 53 colonies during the breeding season, only one colony (Bjørnøya) was not affected, and 12 colonies were not monitored (Figure 1). Positive H5N1 samples were associated with 24 of the 41 sampled colonies (58%), either through direct sampling of dead gannets from the colony or by proximity of dead gannets to colonies. A small colony at Store Ulvøyholmen, Norway (330 Apparently Occupied Nests (AONs) in 2015, Barrett et al. 2017) was reported abandoned (Børge Moe, pers comm) and, since dead gannets were reported close to the colony, this may have been due to HPAI. One gannet sample from a bird found dead at Kjelmøya (Norway) tested positive for H5N5. The earliest outbreaks occurred in the northeast Atlantic in Iceland (at Eldey, Brandur and Raudinupur during 15^th^, 17^th^ and 26^th^ April), followed by Shetland, Scotland (Noss and Hermaness on 1^st^ and 4^th^ May, respectively) then the Outer Hebrides, Scotland (St Kilda, 10^th^ May). Subsequent outbreaks appeared to occur from early June in southern Norway (Runde, 8^th^ June). The concurrent, southwards progression occurred along the east coast of the UK (e.g., Troup Head, 20^th^ May, Bass Rock 4^th^ June). By mid-June, there were HPAIV outbreaks in northern Norway (Syltefjord, 16^th^ June), the southern North Sea (Heligoland, 21^st^ June), the Channel Islands (Les Etacs and Ortac, 28^th^ June) and the southernmost colony Rouzic, France (1^st^ July). In July and early August, signs of HPAI appeared in northwest Norway, the Faroe Islands (Mykineshólmur, 7^th^ July) and in a clockwise progression around the UK, followed by Wales (Grassholm, 21^st^ July) and then in Ireland (Clare Island, Lambay, Bull Rock, Little Skellig, Great Saltee, Ireland’s Eye; 10^th^, 25^th^, 26^th^, 31^st^ August, 1^st^ and 12^th^ September respectively). The northernmost colony Bjørnøya (52 AON in 2016, Barrett et al. 2017) appeared unaffected by HPAIV. No information was available for several remote colonies in the west and northwest of Scotland but unusually high mortality at Sule Skerry was detected after the breeding season in October (Wanless and Harris, in press).

**Figure 1:**
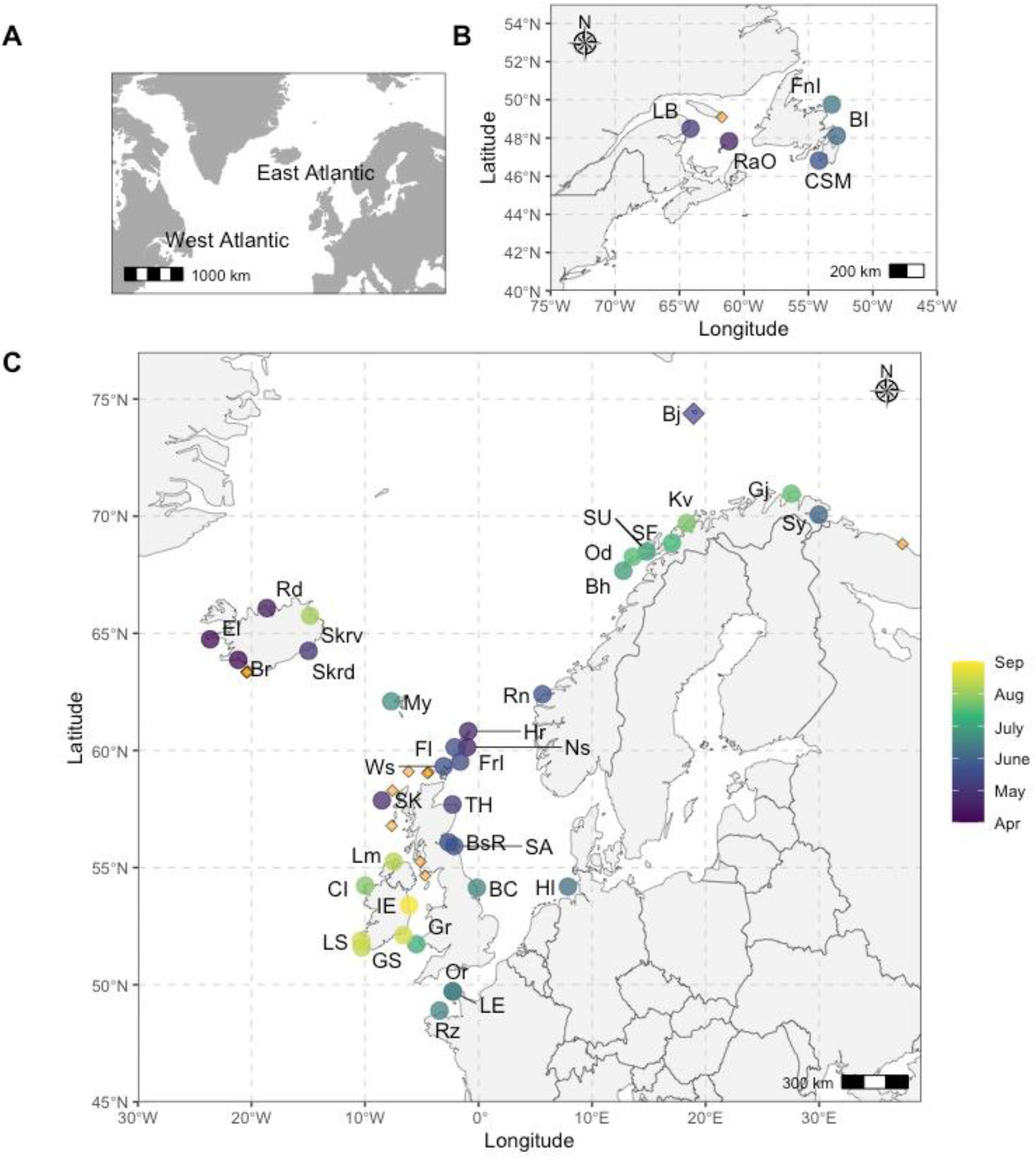
The timing of HPAIV outbreaks across the gannet metapopulation in 2022, based on the first date unusual mortalities in adults were observed. Affected colonies (n = 40) are indicated by circles, coloured by date. Colonies where information is unavailable (n=12) are indicated by orange diamonds. Letter combinations indicate colony name abbreviations, for full colony name see Supplementary Online Material, Table S1). A) Geographical context. B) Colonies in the West Atlantic. C) Colonies in the East Atlantic. A navy-coloured diamond indicates Bjørnøya (Bj, Norway, the northernmost colony, H Strøm pers. obs.) where no signs of HPAIV were observed. The Store Ulvøyholmen colony (SU) was found abandoned (confirmation received 29th June 2022 in litt.) No signs of HPAIV were detected in the colony Ailsa Craig (AC) between Northern Ireland and Scotland on the 28^th^ July 2022, but there was no visit later in the season when the surrounding colonies were affected.

The outbreaks in the northwest Atlantic metapopulation developed in parallel to these in the northeast, with the earliest outbreaks occurring between early and mid-May in the three colonies in the Gulf of St. Lawrence (at Rochers aux Oiseaux, Magdalen Islands, 1^st^ May and, Le Bonaventure, 20^th^ May) followed by the colonies in Newfoundland throughout June (Cape St. Mary’s, 6^th^ June, Baccalieu Island, 17^th^ June, and Funk Island, 24^th^ June).

### b. Case Study: Impact of HPAIV on Bass Rock

Unusually high gannet mortality during incubation in early June 2022 was the first suggestion of an HPAIV outbreak at the Bass Rock and subsequent testing of four carcasses from 4^th^ June proved positive for clade 2.3.4.4b HPAIV H5N1.

#### I. Impact of HPAIV on apparently occupied sites, breeding success and adult survival

##### Apparently occupied sites

A total of 21,227 live birds were counted on the 30^th^ June 2022. An additional 5,035 birds were identified as dead, approximately 3.3% of the breeding population (assuming 150,518 breeding adults from 75,259 AOS, Murray et al. 2015), however, many additional birds will have died at sea. Given the almost complete absence of immatures and non-breeders at the colony during June, it is highly likely that the majority of birds counted, both live and dead, would have been breeding adults.

##### Breeding success

Monitored nests declined from 93 to 23 (75% decline) between 15^th^ June and 14^th^ August. However, empty nest sites on the 15^th^ June indicated nests had already failed prior to the start of monitoring (Figure 2). The majority of the 93 nests had failed by the beginning of July with nest abandonment leaving gaps within the colony (Figures 2 and 3). An index of breeding success was estimated as 0.247 based on the presence of 23 large, apparently healthy chicks in the study areas on the 14^th^ August. Clinical signs of viral infection, seizures, and lethargy were observed in a small number of chicks (aged 2+ weeks) outside of our study areas, but since they were not monitored their fate is unknown.

**Figure 2:**
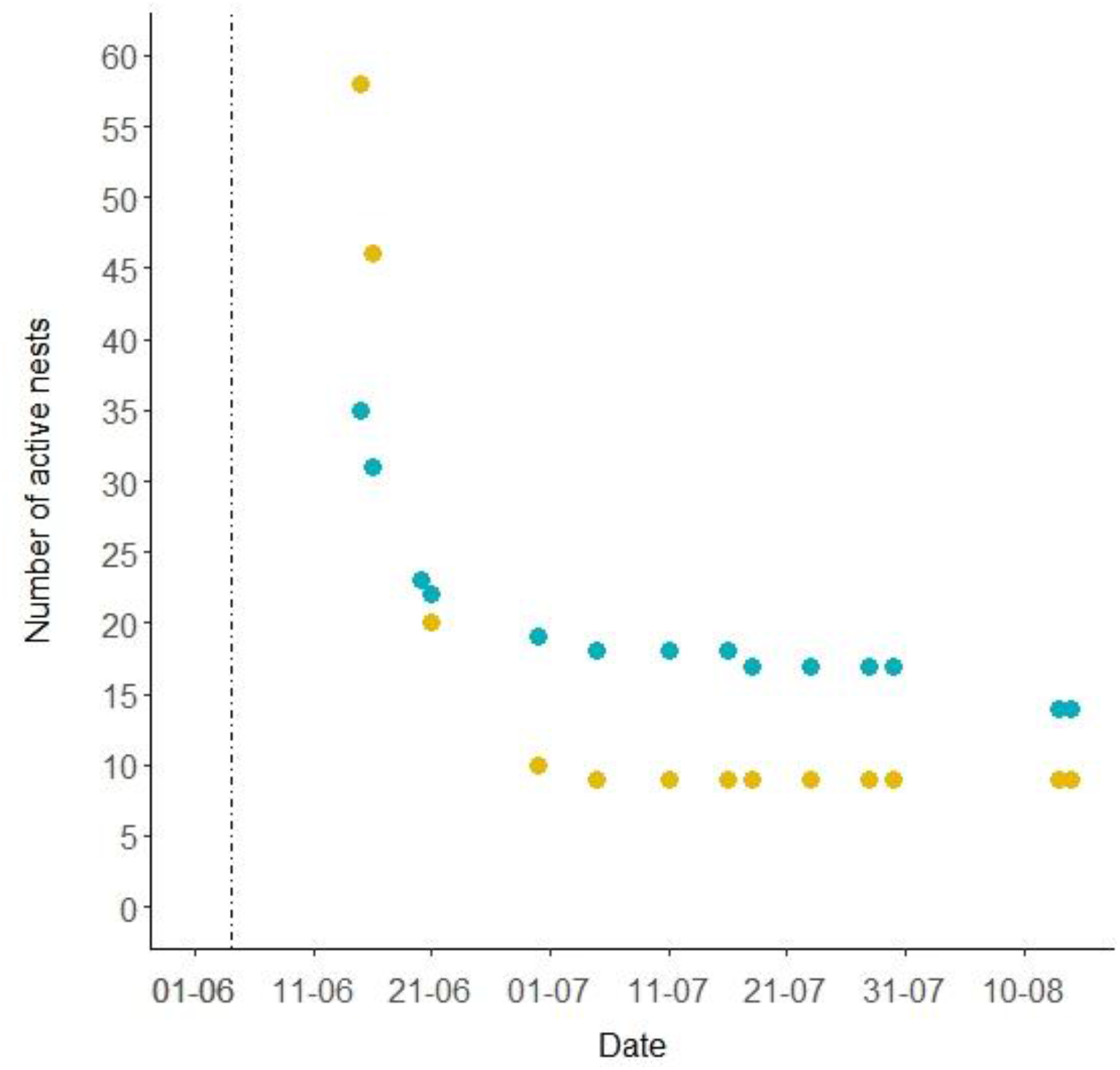
The number of active nests within two study areas on Bass Rock; area 1 in blue, area 2 in yellow. Dotted vertical line indicates 4^th^ June, the date carcasses were collected for testing by the Animal Plant Health Agency (APHA).

**Figure 3.**
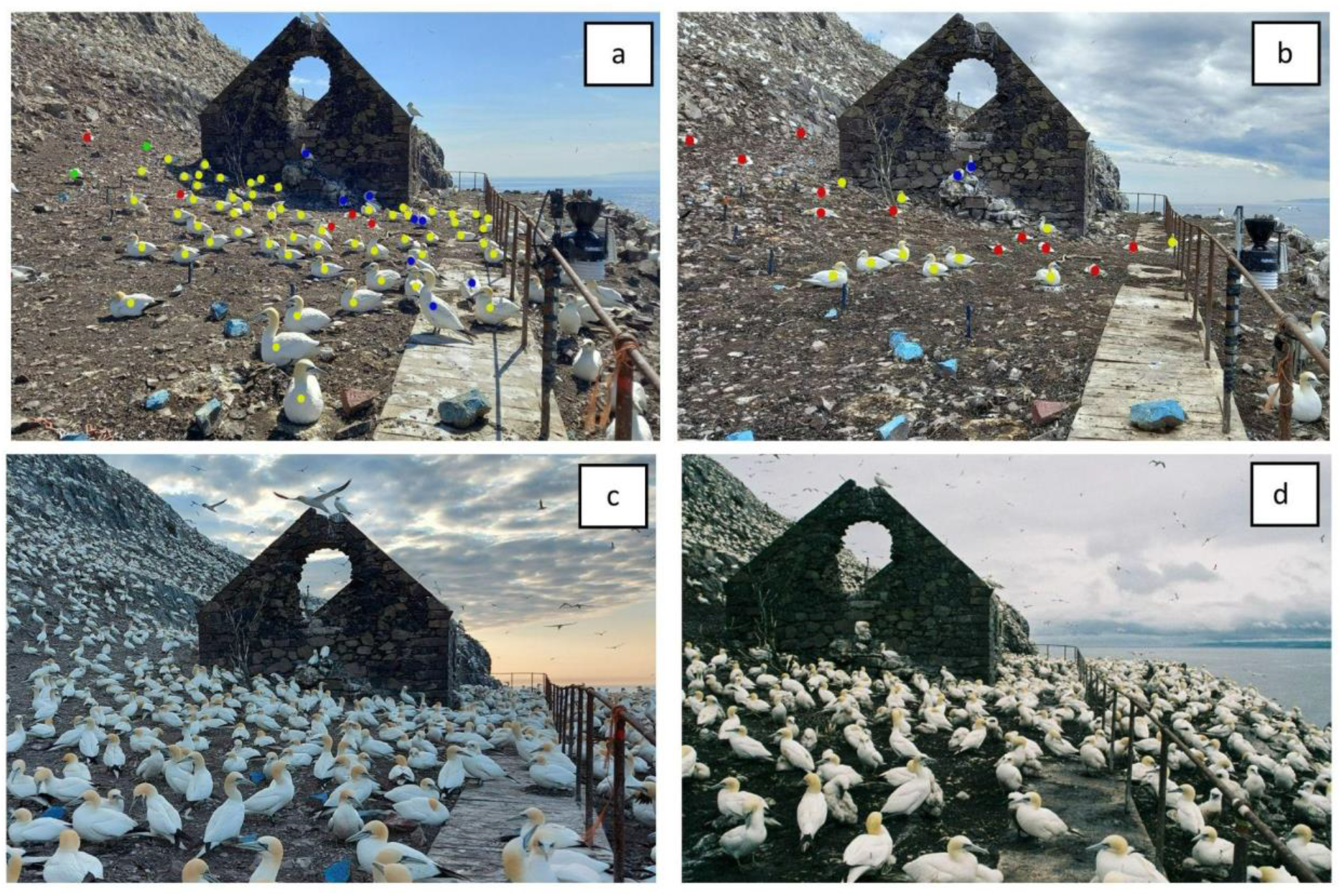
Nest failures, apparent from gaps between birds, and dead birds in study area 2 of the Bass Rock on **(a)** 15^th^ June and **(b)** 30^th^ June, **(c and d)** study area on 29^th^ April 2022 and 23^rd^ July 2020 showing typical nest spacings and densities for the respective time of year; **(c)** pre-laying and **(d)** mid-chick rearing. Coloured dots in **a)** and **b)** indicate the status of the bird; red - dead, green - visibly sick, yellow - active nest with healthy adults, dark blue - non-breeding healthy adults, light blue painted rocks delineate a path.

##### Adult survival

The top model showed strong support for survival probability varying with time and for re-sightings to vary with time following the first year after marking (Table 1). Adult survival between 2021 and 2022 was 0.455 (95% CI: 0.153 – 0.794) compared with an average annual survival of 0.940 (SD 0.035) between 2011 and 2021. The resighting probability during 2022 was 0.615 (95% CI: 0.144 – 0.938) compared with an average of 0.839 (SD: 0.066) between 2011 and 2021.

**Table 1.** Candidate model set for estimating annual survival of northern gannets from Bass Rock between 2010 and 2022. Inflation factor (c) = 1.212. Effects fitted to apparent survival (ϕ) and resighting probabilities (*p*) (*t*: time dependent; *c*: time constant). AICc: Akaike Information Criterion for small samples. ΔAICc: difference in AICc between model in question and best model. Num. Par.: number of parameters.

| Model | QAICc | ΔQAICc | AICc<br>Weights | Model<br>Likelihood | Num. Par. | QDeviance |
| --- | --- | --- | --- | --- | --- | --- |
| $\phi(t) p(c/t)$ | 2031.27 | 0.00 | 0.912 | 1.000 | 24 | 377.31 |
| $\phi(c) p(c/t)$ | 2036.85 | 5.58 | 0.056 | 0.061 | 13 | 405.39 |
| $\phi(t) p(c/c)$ | 2037.94 | 6.67 | 0.032 | 0.036 | 14 | 404.45 |
| $\phi(c) p(c/c)$ | 2225.89 | 194.62 | 0.000 | 0.000 | 3 | 614.63 |

Seven colour-ringed birds were found dead during June and July 2022 on the North Sea coasts of the UK, Sweden and Denmark, and eight were found dead on the colony in October, compared with 3 dead recoveries between 2015 and 2021.

#### II. Serology and iris colour

All 18 birds tested negative for viral nucleic acid from cloacal swabs, indicating they were not currently infected. Of the 19 blood samples, two were insufficient for testing and eight tested positive for H5 antibodies indicating a previous infection.

Black irises – instead of the usual pale blue – were first noted on 15-16^th^ June 2022. Iris colour varied from completely black to mottled and with some variation between eyes and did not present like a dilated pupil (Figure 4). The likelihood of testing positive for HPAIV H5 antibodies was higher in birds with black irises (77.7%) compared to birds with normally coloured eyes (12.5%; Fisher’s exact test; p <0.05). The hemagglutinin (HA) binding antibody levels in serum samples, as detected by a Haemagglutination Inhibition (HAI) titre, were 1/16 (n = 3) and 1/32 (n = 5, including the sample from the bird with healthy irises) (Table 2 and Supplementary Online Material, Table S2).

**Figure 4.**
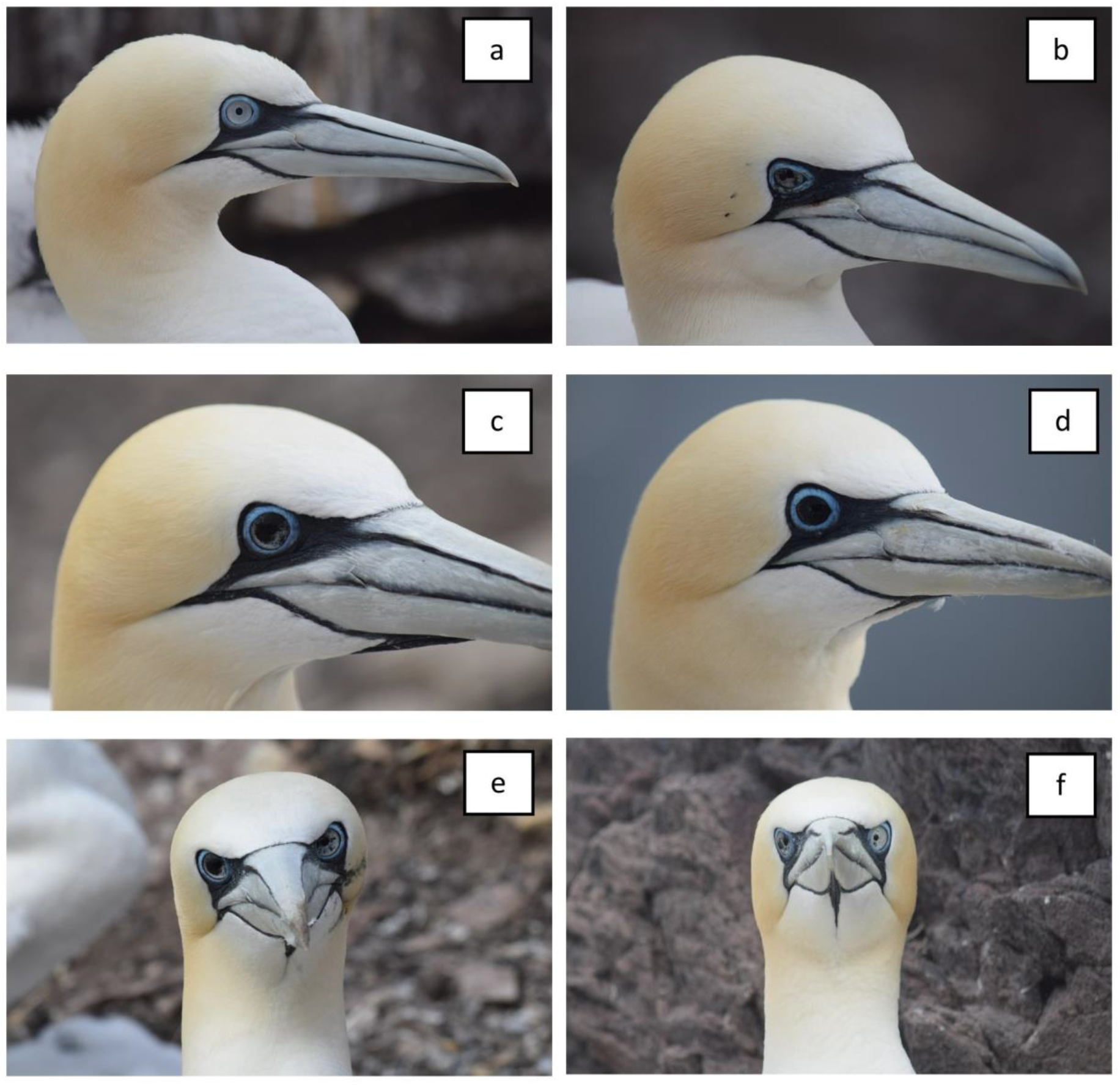
Gannets on the Bass Rock colony in 2022 with black flecking in their irises. The condition was variable between individuals from **a)** healthy, **b** and **c)** increasing degrees of black flecks in the iris **d)** completely black iris, and asymmetrical irises affected to **e)** greater and **f)** lesser extents. No pattern was detected in the asymmetry of black irises.

**Table 2.**
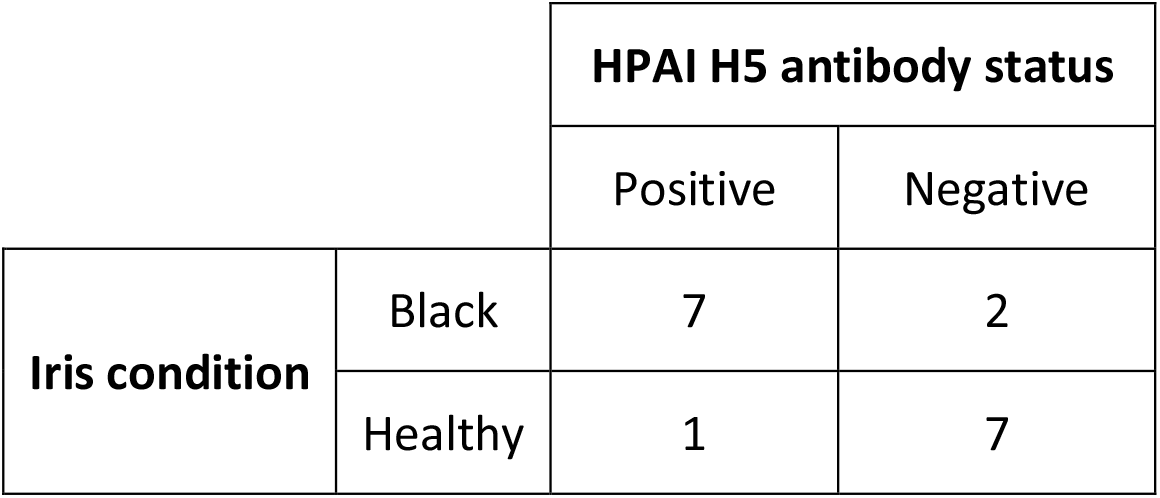
Serological results from 17 adult gannets from Bass Rock tested for H5 antigen.

|  |  | HPAI H5 antibody status |  |
| --- | --- | --- | --- |
|  |  | Positive | Negative |
| Iris condition | Black | 7 | 2 |
|  | Healthy | 1 | 7 |

## 4. Discussion

### a. Global Context: HPAIV spread across the North Atlantic gannet metapopulation

During summer 2022 HPAIV H5N1 was recorded for the first time in gannets, causing mortality on an unprecedented scale across their entire Atlantic breeding range. Positive tests from 58% of monitored colonies mean it is likely that unusually high mortality in the 16 untested colonies in 2022 was due to HPAI. Of the 41 colonies monitored, only one was confirmed to have been unimpacted/unaffected. Strong evidence of an HPAIV outbreak at a colony unmonitored during the breeding season, Sule Skerry, northern Scotland (Harris and Wanless in press) suggests it is likely some of the 12 remote unmonitored colonies were also affected.

All positive samples collated across the northeast and northwest Atlantic metapopulations were subtype H5N1 apart from a single gannet sample testing positive for subtype H5N5 from the Sør-Varanger municipality in Troms and Finnmark county, Norway. In Norway, Subtype H5N5 has also been detected in 30 birds from different species, including White-tailed Eagles, gulls (*Laridae*), Great Skuas and corvids (*Corvidae*) (S. Granstad, personal communication, March 26, 2023).

Although a thorough estimation of gannet mortality during the 2022 HPAIV outbreak is beyond the scope of the paper, we document the spread of the HPAIV outbreak and provide details on impacts at the largest gannet colony. Following the first confirmed cases in Iceland during April 2022, HPAIV was detected almost simultaneously across the northeast and northwest Atlantic metapopulations. HPAIV outbreaks, confirmed and inferred from dead untested birds, occurred in at least 75% of all 53 known gannet colonies. While gannets are a well-studied species, we note that sampling effort was not standardised among colonies (e.g. uncertainty in data from northern Norway, Iceland and some of the Irish colonies is largely due to the use of passive surveillance data rather than direct colony monitoring), but we have no reason to believe this leads to an inaccurate representation of the timing of HPAIV outbreaks.

### b. Possible mechanism of HPAIV transmission between gannet colonies

The scale and speed at which HPAIV spread through the gannet metapopulation was dramatic, but the mechanism of transmission and the subsequent spread between colonies is unclear. A possible source may have been infectious gannets returning from their wintering areas. During the spring migration, gannets in the eastern North Atlantic frequently perform a clockwise loop around the UK, with Icelandic breeders arriving earlier than those breeding on the Bass Rock (Furness et al. 2018). However, gannets from different colonies overlap to some degree in the wintering areas (Fort et al. 2012; Furness et al. 2018), making the sequential nature of the spread less likely due to differences in migratory timing. Yet an unprecedented stranding of dead adult gannets on the Dutch coast prior to the start of the breeding season in April 2022 potentially indicates HPAIV exposure over the previous winter although none of these birds were tested (Camphuysen et al. 2023).

The timing of outbreaks on each side of the Atlantic and throughout the northeast metapopulation, might point towards HPAIV transmission via other infected seabirds. Great Skuas (Grecian et al. 2016) were severely affected by HPAIV H5N1 in Scotland in 2021 (Banyard et al. 2022) and again in 2022 (Camphuysen et al. 2022, Falchieri et al. 2022). Great Skua breed in close proximity to gannets in Iceland, the Faroes and northern Scotland (Birdlife International, 2023) and overlap with the winter range of gannets from both sides of the North Atlantic (Magnusdottir et al. 2012, Fifield et al. 2014). Great Skua regularly kleptoparasitise gannets (Anderson 1976) which in addition to transmission via faeces and respiratory secretions could explain the spread across taxa. Brown Skuas (*Stercocarius antarcticus)* are likely vectors of avian cholera on Amsterdam Island, Indian Ocean (Gamble et al. 2019) and we speculate a similar role for Great Skuas triggering the HPAIV outbreak in gannets in 2022. Yet this does not explain the subsequent spread through the gannet metapopulation, and questions remain about why spill-over into gannets may or may not have occurred during the 2021 outbreak among skuas. Similarly, waterfowl and gull species have been found to play an important role in intercontinental transmission of LP and HPAIVs via Iceland, the link between the East Atlantic and North American Atlantic Flyways (Duesk et al. 2014). Gulls are known to frequent seabird colonies to opportunistically prey on eggs and chicks (Donehower et al. 2007, pers obs) and may therefore have played a role in virus spread.

The subsequent clockwise spread around the UK seems unlikely to be linked to centrally-placed adults foraging at sea, based on current evidence. During chick-rearing, gannets have colony-specific foraging ranges with limited overlap (Wakefield et al. 2013) and tend to have individual specific foraging grounds (Wakefield et al. 2015, Votier et al. 2017). However, the HPAIV outbreak may have altered their movement behaviour leading to an increased inter-colony contact (Jeglinski et al. in prep; d’Entremont and Montevecchi unpubl. data). Immature gannets are another possible route for spreading the virus while prospecting among colonies (Votier et al. 2011). They also have larger foraging ranges than breeders (Votier et al. 2017, Grecian et al. 2018), and therefore a greater chance of inter-colony overlap. Nevertheless, immature gannets tend to return to the colony much later than adults, being scarce during April-May and only appearing in large numbers during June/July (Wanless 1983, Nelson 2002), so were unlikely to have played a role during outbreaks during April and May, though they may have played a role during outbreaks later in the breeding season (Figure 1). More research into virus incubation and length of infectious period in addition to possible transmission pathways between species that overlap in their wintering, migratory and breeding areas is paramount (Hill et al. 2022).

### c. Case Study: Impact on Bass Rock Gannet Colony

Drone footage on 30^th^ June recorded 5,035 dead individuals that represented ∼3% of breeding adult gannets on Bass Rock. This is likely an underestimate as it excludes decomposed birds or those that died at sea and does not account for the colony growth since 2014 (Murray et al. 2017). This figure compares with an estimated 7% mortality at Mykineshólmur, Faores (unpublished) and 6% at Sule Skerry, Scotland (Harris and Wanless, in press), both from aerial counts although the Sule Skerry count was performed at the end of the breeding season. Drone counts in late June indicate that the colony was ∼71% smaller than during the last full colony count in 2014 (Murray et al. 2015). However, the colony had grown since 2014 (Murray 2017) so again, this is almost certainly an underestimate, though note that different methodologies and counting units make a direct comparison difficult.

Around one quarter of nests with an egg on 15^th^ June still had a chick in late August, which is much lower than the mean UK gannet breeding success during 1961–2018 (mean ± standard deviation) 0.72 ± 0.12 (Jeglinski et al. 2023). There are methodological differences in approach, but the comparison provides a further indication of the severe impact of the virus. The primary cause of breeding failure appeared to be nest abandonment, either when adults did not return from foraging trips or died at the nest.

Adult survival was approximately 42% lower than the average of 0.940 (SD 0.035) between 2011 and 2021. The reduction in the number of re-sighted colour-ringed birds indicates that a large proportion of adults have died, but a full assessment of the impact on adult survival will have to wait until 2023 when visual searches will be made for returning birds. Similar to most seabirds, gannets are a long-lived species making their populations particularly sensitive to changes in adult survival therefore the consequences of a significant reduction in adult survival could be considerable (Croxall & Rothbury, 1991).

Despite a modest sample size, our study suggests that gannets infected with HPAIV H5N1 can survive, with important implications for the long-term consequence of the virus impact. We also found that black iris coloration in otherwise apparently healthy gannets is a likely indicator of prior infection. One seropositive bird had healthy irises, but this may be related to a different subtype of HPAIV or LPAIV (Wilson et al. 2013), to waning antibody levels following prior infection, or may suggest that not all infected birds develop black irises. We suggest the two birds with black irises that tested negative for antibodies had previously been infected but had already lost the antibodies, however further investigation is needed to inform on antibody persistence. Black eyes have been reported in gannets once before, but the reason is unknown (J. Swales pers comm., Balfour 1922). During the HPAIV outbreak in 2022, gannets with black irises were also reported from colonies in the UK (Bempton Cliffs, Grassholm and Ortac), France (Rouzic), Germany (Heligoland) and Canada (île Bonaventure). In early spring 2023, gannets with black irises were observed at the Bempton Cliffs, Bass Rock, Troup Head, Rouzic and Les Etacs colonies, suggesting the potential for a longer-lasting or even permanent modification of the iris.

## 5. Recommendations

Future research should quantify changes in demography (i.e. population size, adult survival and breeding success) of gannets and other impacted seabirds while also assessing whether previously infected birds have developed immunity in order to model disease progression and long-term impacts of HPAIV (Hill et al. 2019). Additionally, assessments of infection and mortality rates in different age classes, and of how previous infection might influence fertility or the outcome of a second infection are also needed (Wilson et al. 2013). Juvenile gannets have been found to carry antibodies to HPAIV (Grémillet et al. in prep) but it is unknown whether these were maternally derived or produced in response to infection (DeVriese et al. 2010).

Black irises may provide a useful non-invasive diagnostic tool, more work is required to better understand its efficacy, if it applies to any other species, and whether there are any potential costs in terms of vision. Ophthalmology exams or histopathology examinations are also required to determine what is causing the black colouration. It is also desirable to better understand the circulation of LPAIVs and prior exposure to antigenically related HPAIV sub-types in seabird populations to better understand potential cross-protective immunity, as well as the potential for compensatory recruitment to offset mortality (Votier et al. 2008, Jeglinski et al. 2023).

If sampling for live virus, we recommend cloacal swabs be taken in conjunction with oropharyngeal swabs (Suarez et al. 2000, van den Brand et al. 2018) because of possible differences in virus genotype detectability (Slomka et al, in prep). Primary flight feathers can also be used as a diagnostic indication of systemic viral infection as infectious virus can be detected in these samples (Nuradji et al. 2015). The 2022 HPAIV H5N1 outbreak has provided another significant stressor to those already faced by our rapidly declining seabird populations (Dias et al. 2019, Careen et al. 2023) - quantifying and perhaps even mitigating its impact is therefore crucial if we hope to see a healthy seabird assemblage across the world’s oceans.

## Acknowledgements

This work was funded by the Forth and Tay Offshore Wind Farm Developers; Neart na Gaoithe Offshore Wind Ltd, Seagreen Wind Energy Ltd and SSE Renewables, the Animal Plant Health Agency, and the UK Department for Business, Energy and Industrial Strategy Offshore Energy Strategic Environmental Assessment BEIS OESEA programme, as well as a NERC Urgency Grant NE/X013502/1 and Natural Sciences and Engineering Research Council of Canada (NSERCC) and Fisheries and Oceans Canada. ACB, JJ, TL and IHB were part supported by the Biotechnology and Biological Sciences Research Council (BBSRC) and Department for Environment, Food and Rural Affairs (Defra, UK) research initiative ‘FluMAP’ (grant number BB/X006204/1). Funding was also provided by the Defra and the Devolved Administrations of Scotland and Wales, through grants SE2213 and SV3006. UAV data collection was undertaken by the University of Edinburgh’s Airborne Research and Innovation facility (ARI), the NERC Field Spectroscopy Facility (FSF)) and Scotland’s Rural College (SRUC), in partnership with the Scottish Seabird Centre.

We thank Sir Hew Hamilton-Dalrymple and the Scottish Seabird Centre, North Berwick for support and access to Bass Rock, and Jack Dale and John McCarter for logistic support. We thank Esbern í Eyðanstovu for, for making the observations of Mykineshólmur available to us. We thank Armel Deniau, Grgoire Delavaud, Aurlien Prudor, Timothe Poupart and Gauthier Poiriez for support with data collection at Rouzic. We thank Andrew Lang, Kathryn Hargan (Memorial University of Newfoundland and Labrador, Canada), Pauline Martigny (Université du Québec à Rimouski) and Ishraq Rahman and Jordan Wright (Memorial University of Newfoundland) for field and lab support. Permission to undertake work on the Bass Rock during the HPAI outbreak was granted by NatureScot. Birds were ringed and swabbed with permits and approval from the British Trust for Ornithology. Blood sampling was carried out under licence from the UK Home Office; Project licence number PEAE7342F and Personal licence number IF2464041.

