## Supplementary Online Material for "High pathogenicity avian influenza (H5N1) in Northern Gannets: Global spread, clinical signs, and demographic consequences"

### S1. Health and Safety, and Biosecurity Protocols – working on Bass Rock

To ensure the safety of birds and field workers, we followed strict biosecurity and health and safety measures that were developed in collaboration with the RSPB, NatureScot, experts at the University of Glasgow and the Animal Plant Health Agency (APHA).

We took every effort to ensure infected material was not transferred from sick to healthy birds or from Bass Rock to the mainland.

For visual monitoring, we wore work Type 5/6 coveralls and, since the colony becomes very dry in the summer with airborne dust/feathers, we wore face masks. At the time bird handling took place, there had been no sign of a bird displaying any clinical signs at the colony for 42 days; however, the presence of subclinical individuals could not be excluded and therefore full PPE was worn.

For bird handling, we used sealable plastic boxes to store and transport sterile clean Personal Protective Equipment (PPE) and used PPE (in a bin liner), respectively. Each field team member wore Type 5/6 coveralls, FFP3 face mask, non-latex gloves and safety goggles that seal around the face. In addition, to limit the risk of cross-infecting birds, we wore a sterile long-sleeved disposable apron on top of coveralls, and we replaced aprons and gloves between handling different birds. The person catching the bird additionally wore cut-resistant arm guards under their coveralls and apron, and cut-resistant gloves under non-latex gloves to ensure there was no possibility of a bite making contact with the skin.

We disinfected all items of equipment with Safe4 disinfectant before handling a different bird. Excellent hand sanitisation was adhered to at all times, Safe4 wipes and hand sanitizer were used at regular intervals and at all times before eating and drinking.

Before leaving the colony, we sealed all used PPE in two layers of black bin liners. Footwear was thoroughly disinfected with a Safe4 footbath, a wire brush used to remove any organic material. Once on the mainland PPE was disposed of in industrial waste.

All samples taken for AIV testing were sent to APHA-Weybridge and tested within a Specified Animal Pathogen Order (SAPO) level 4 and Advisory Committee for Dangerous Pathogens (ACDP) level 3 rated containment envelope using UKAS accredited diagnostic tests.

### S2. Collection of Unmanned Aircraft System data

The Bass Rock was surveyed on the 30th June 2022 by the University of Edinburgh's Airborne Research and Innovation facility (ARI), the NERC Field Spectroscopy Facility (FSF)) and Scotland's Rural College (SRUC), in partnership with the Scottish Seabird Centre. The survey utilised ARI's DJI Matrice 300 RTK unmanned aircraft system fitted with a DJI-Zenmuse L1 LiDAR and photogrammetry sensor. All flights were conducted from the helipad at the southern tip of the Bass Rock with the system's Real-Time-Kinematic (RTK) base station set up on a marked point adjacent to the pad. The RTK system ensured that GNSS positional corrections were transmitted in real time to the aircraft and on to the sensor, providing cm-level precision across the dataset with respect to the base station location. Future surveys will therefore be repeatable with cm-level precision.

The topography of the island slopes steeply upwards from the south to the north, so in order to stay within the legal maximum of 400ft (120m) from the nearest point on the surface, separate mission plans were designed to effectively contour up the island in steps. This also allowed the crew to ensure that continuous line of sight was maintained with the aircraft throughout the operation, as required by law. Operating at this height yielded a ground sampling distance (GSD) of approximately 3cm across the island. Several oblique images were also acquired from the southern, eastern and western sides of the island, but this was not possible from the north due to line-of-sight limitations.

During an initial test flight when the aircraft was launched upwards from the helipad, gannets in flight above the colony showed apparent interest in the aircraft. This interest occurred at a height of ~50-60m above the colony where the aircraft and gannets arriving and departing the colony occupied the same height range. The interest in the aircraft was evident by birds flying towards the aircraft, apparently to investigate, sometimes in large numbers. No contact was made between birds and the aircraft, and no aggressive behaviour was observed.

To avoid disturbance to gannets on the colony and in flight above and around the colony, the aircraft was relaunched and, following a brief control check, flown at $4ms^{-1}$ and approximately 10m above the sea to a range of 200-250m offshore, beyond most gannets in flight. The aircraft was ascended as rapidly as possible to a height of 100m (above flight height) before bringing it back in towards the island. At a height of 100m above the island, the aircraft was flying above the gannets in flight, and they showed no interest in the aircraft. The same flying strategy was used in reverse prior to landing with the aircraft flown out to 200-250m offshore at a height of 100m before being descended to 10m above sea level and flown back into the helipad.

The resulting imagery was processed through Agisoft Metashape (Agisoft LLC, St Petersburg, Russia) software to produce an orthomosaic of the entire Bass Rock with GSD of approximately 3cm.

**Table S1: Names, abbreviations and the date of first appearance of HPAIV outbreaks based on unusually high mortality in adult gannets for all colonies for which we were able to collate such data (n = 40). Colonies are ordered by the date on which HPAIV was first recorded. HPAIV test results for colonies (n = 24) are stated as positive if dead gannets were collected for testing from a colony or the geographic origin of a positive gannet sample was associated with the nearest gannet colony - note that the sampling date does not coincide with the first date of HPAIV outbreak.**

| | Colony (abbreviation) | Country | First date of HPAIV outbreak | HPAIV test results |
|---|---|---|---|---|
| 1 | Eldey (El) | Iceland | 15-04-2022 | positive |
| 2 | Brandur (Br) | Iceland | 17-04-2022 | positive |
| 3 | Raudinupur (Rd) | Iceland | 26-04-2022 | positive |
| 4 | Noss (Ns) | UK | 01-05-2022 | positive |
| 5 | Rochers aux Oiseaux, Magdalen Island (RaO) | Canada | 01-05-2022 | NA |
| 6 | Hermaness (Hr) | UK | 04-05-2022 | positive |
| 7 | St. Kilda (SK) | UK | 10-05-2022 | NA |
| 8 | Skrudur (Skrd) | Iceland | 12-05-2022 | positive |
| 9 | Troup Head (TH) | UK | 20-05-2022 | NA |
| 10 | Le Bonaventure (Bn) | Canada | 20-05-2022 | positive |
| 11 | Fair Isle (FrI) | UK | 30-05-2022 | NA |
| 12 | Foula (Fl) | UK | 01-06-2022 | NA |
| 13 | Westray (Ws) | UK | 03-06-2022 | NA |
| 14 | Bass Rock (BsR) | UK | 04-06-2022 | positive |
| 15 | Cape St. Mary's (CSM) | Canada | 05-06-2022 | positive |
| 16 | St. Abbs (SA) | UK | 05-06-2022 | NA |
| 17 | Runde (Rn) | Norway | 08-06-2022 | positive |
| 18 | Syltefjord (Sy) | Norway | 16-06-2022 | positive |
| 19 | Baccalieu Island (BI) | Canada | 17-06-2022 | NA |
| 20 | Heligoland (Hl) | Germany | 21-06-2022 | positive |
| 21 | Funk Island (FnI) | Canada | 24-06-2022 | positive |

| | | | | |
|---|---|---|---|---|
| 22 | Ortac (Or) | UK | 28-06-2022 | positive |
| 23 | Les Etacs (LE) | UK | 28-06-2022 | positive |
| 24 | Rouzic (Rz) | France | 01-07-2022 | positive |
| 25 | Bempton Cliff (BC) | UK | 02-07-2022 | NA |
| 26 | Mykineshólmur (My) | Faroe Islands | 07-07-2022 | NA* |
| 27 | Buholmene (Bh) | Norway | 13-07-2022 | positive |
| 28 | St. Ulvøyholmen (SU) | Norway | 16-07-2022 | NA |
| 29 | Grassholm (Gr) | UK | 21-07-2022 | positive |
| 30 | St. Forøya (SF) | Norway | 27-07-2022 | NA |
| 31 | Oddskjæren (Od) | Norway | 31-07-2022 | positive |
| 32 | Gjesvær (Gj) | Norway | 02-08-2022 | NA |
| 33 | Kvitvær (Kv) | Norway | 08-08-2022 | positive |
| 34 | Clare Island (CI) | Ireland | 10-08-2022 | NA |
| 35 | Skoruvikbjarg (Skrv) | Iceland | 19-08-2022 | NA |
| 36 | Lambay (Lm) | Ireland | 25-08-2022 | positive |
| 37 | Bull Rock (BlR) | Ireland | 26-08-2022 | positive |
| 38 | Little Skellig (LS)** | Ireland | 31-08-2022 | NA |
| 39 | Great Saltee (GS) | Ireland | 01-09-2022 | positive |
| 40 | Ireland's Eye (IE) | Ireland | 12-09-2022 | positive |

* Four gannets found on the Faroe Islands (distance to Mykineshólmur between ~ 20 - 50 km) tested positive for HPAIV in early June, but the first signs of abnormal mortality at the colony were only observed about a month later. We therefore did not associate the positive samples with the Mykineshólmur colony.

** Positive samples were retrieved close to Little Skellig on the 10th August 2022, but the first signs of unusually high mortality at the colony were only observed three weeks later. We therefore did not associate the positive samples with the Little Skellig colony.

**Table S2. Serology results for H5 antibodies and iris condition of 17 gannets on Bass Rock in September 2022.**

| Sample number | Bird ID | Breeding status | Right eye | Left eye | HAIT Titre | Interpretation |
|---|---|---|---|---|---|---|
| 1 | 1491466 | On territory | 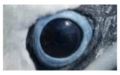 | 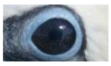 | 1/16 | **H5 +ve** |
| 2 | 1491467 | On territory | 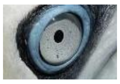 | 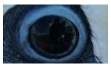 | 1/16 | **H5 +ve** |
| 3 | 1491475 | On territory | 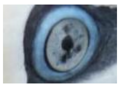 | 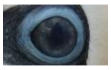 | 1/16 | **H5 +ve** |
| 4 | 1491469 | On territory | 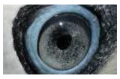 | 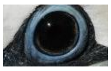 | 1/32 | **H5 +ve** |
| 5 | 1491473 | On territory | 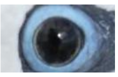 | 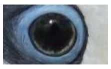 | 1/32 | **H5 +ve** |
| 6 | 1491480 | On territory | 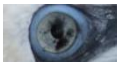 | 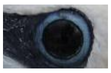 | 1/32 | **H5 +ve** |
| 7 | 1491481 | On territory | 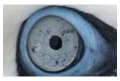 | 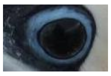 | 1/32 | **H5 +ve** |
| 8 | 1491482 | Chick | 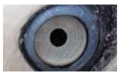 | 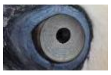 | 1/32 | **H5 +ve** |
| 9 | 1491465 | Chick | 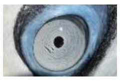 | 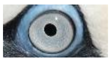 | <1/2 | H5 -ve |
| 10 | 1491468 | Chick | 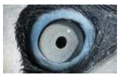 | 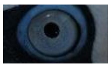 | <1/2 | H5 -ve |
| 11 | 1491470 | Chick | 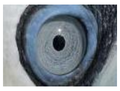 | 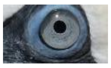 | <1/2 | H5 -ve |
| 12 | 1491471 | Chick | 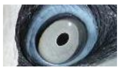 | 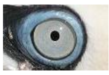 | <1/2 | H5 -ve |
| 13 | 1491472 | Chick | 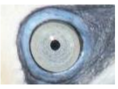 | 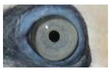 | <1/2 | H5 -ve |
| 14 | 1491474 | Chick | 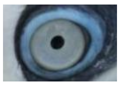 | 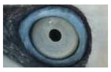 | <1/2 | H5 -ve |
| 15 | 1491476 | On territory | 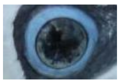 | 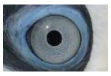 | <1/2 | H5 -ve |

| | | | | | | |
|---|---|---|---|---|---|---|
| 16 | 1491478 | On territory | 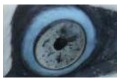 | 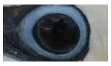 | <1/2 | H5 -ve |
| 17 | 1491479 | On territory | 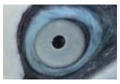 | 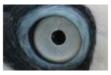 | <1/2 | H5 -ve |

97
